# Comparison of Protein Extraction Methods and Data Analysis Strategies for Complete Metaproteomic Soil Analysis

**DOI:** 10.1101/2024.06.13.598917

**Authors:** Abigale S. Mikolitis, Phillip M. Mach, Marie E. Kroeger, Ethan M. McBride, Trevor G. Glaros

## Abstract

Considerable microbial diversity has been discovered in soil through genomic sequencing. Despite its role in biogeochemical cycling, relatively little is known about the proteomic diversity of the soil microbiome as most commercially available soil kits focus on DNA/RNA extractions. Consequently, a plethora of protein extraction techniques have been developed for soil but have yet to be integrated into simplified, modern sample preparation techniques such as the S-Trap™. Furthermore, classical data analysis strategies for soil metaproteomics rely on genomically-informed databases for peptide/protein identification. This assumes that DNA/RNA extracts adequately represent the soil proteome. Within this study, we systematically assess several extraction techniques, developing a data processing pipeline which is driven by both proteomics and genomics to fully characterize the soil microbiome. Both pipelines reveal remarkably complementary data, with ∼60% of the protein identifications coming from Proteomically-derived databases. Sodium dodecyl sulfate-based extractions proved to provide the most unique protein identifications (∼3000 proteins), and by combining both proteomic and genomic-based results, the total protein identifications increased approximately 2-fold for each extraction. Combining these complementary data pipelines with improved extraction techniques can allow for drastically improved proteomic results (12,307 unique protein identifications), even from minute (50 mg) sample volumes. These enhancements to previous workflows can better describe the microbial diversity within soil and provide a deeper functional understanding of the soil microbiome.

## Introduction

The soil microbiome represents one of the most diverse ecosystems on the planet. Billions of microbes can be present in just one gram of soil^1^, and tens of thousands of species can be active in a given area^2^. Soil microbiome biodiversity, not unlike that of the human gut microbiome, is known to be vast and critical for the health of the local environment. Many microbial functions have been the subject of extensive research, and microbes within soil are known to promote plant growth by cycling nutrients^3^, regulating soil organic matter (SOM)^4^, and producing metabolites useful in both medicine and agriculture^5^; however, many functional aspects of the soil microbiome remain relatively unexplored.

Metaproteomics has emerged as a powerful tool in recent decades to explore the functional dynamics within the soil microbiome apart from its genomic diversity, as achieved by DNA sequencing^6-8^. Metaproteomics involves the identification and/or quantification of the entire set of proteins derived from all microbial activity present within a given environment^9^. In many cases, due to unique peptides, it is possible to attribute some proteomic changes to a particular subset of organisms. This allows for a far richer, albeit more complex, dataset compared to normal proteomics workflows where proteins are extracted from a much simpler, less diverse system (e.g., single cell, cell culture lysate, or tissue/organ). Due to the recent widespread adaptation of high-resolution mass spectrometry as well as significant advancements in data processing algorithms such as de novo peptide sequencing, it is now possible to agnostically interrogate the entire proteome of the soil microbiome.

Given the potential microbial diversity within an area of a given soil type, the theoretical proteome database search space is practically infinite. This computational problem is presently insurmountable even with scalable cloud-based computing resources. Given the potential reference database size, even with an acceptable false discovery rate (<1%), there would be thousands of false peptide/protein identifications which would wholly convolute the data. As such, microbial-focused metaproteomic studies have largely been predicated on advances made by soil metagenomics^10, 11^. Using the microbial diversity established from metagenomics, a finite number of representative organisms can be inferred to limit the proteomic search space allowing for a superficial look at the proteome within a given sample. This strategy implies that the extraction techniques used for DNA/RNA are in fact representative of the protein content accessed using a completely different extraction technique. Despite this assumption, metaproteomic analyses have already been performed successfully for other difficult matrices including activated sludge from wastewater treatment^12^, ocean water^13^, and the human gut^14^.

The first and arguably most crucial step in soil metaproteomics is sample preparation. Protein extraction from soil has been shown to require special considerations to ensure sample integrity; porous microaggregates and variability in soil composition often limit the number of proteins extracted15, 16. Of the available extraction methods currently within the literature, significant variability exists in terms of efficiency 16. Variables such as pH, organic matter content, and the presence of humic substances can create not only difficulty in extracting proteins, but also results in protein modification prior to analysis^17^. Further complicating the issue, there are no commercially available protein extractions kits for soil, unlike DNA and RNA, which has led to a plethora of available techniques with very little evidence available to systematically compare one to another.

One commercially available kit for the extraction of proteins is known as the S-Trap™ (Protifi, LLC, Farmingdale, NY). This kit, which uses a streamlined extraction method similar to filter aided sample preparation (FASP), is able to extract proteins using a simple spin column. This spin column, either in single or 96-well format, can be used to solubilize, reduce, alkylate, trap, clean, and digest proteins all from a single substrate. Buffers can be purchased or made separately for each step, and different kits are available for different protein concentrations within a sample. Although these kits have been used successfully for extracting proteins from a variety of different matrices including sputum, urine, and whole cell lysates^18-20^, to date they have not been evaluated for the extraction of proteins from soil. To compare the utility of the S-Trap™ with several extraction methods, we have performed a systematic evaluation of the S-Trap™ utilizing common soil extraction methods for the metaproteomic analysis of soil extracts. These extraction methods are compared in terms of protein extraction efficiency, and protein abundance reproducibility was evaluated for the best extraction type with several different masses of soil. The proteomics data sets were then compared with traditional metagenomic data to determine heterogeneity between the proteomic and genomic data sets.

## Materials and Methods

### Soil Analyses

Arid (sandy-type) soil was collected from the area within Los Alamos National Laboratory. This Penistaja-like series soil type can be characterized as sandy loam close to the surface, as is typical for this area within New Mexico. The top 2 cm of surface soil was collected using 50 mL conical tubes, and five tubes were collected in total from various areas of the site. All five replicates of the arid sandy-type soil collected were homogenized into one sample. The arid soil was sieved with a 2 mm mesh sieve and stored at -20 °C until processing.

Total soil organic carbon (SOC) measurements were completed by the Geology and Geochemistry Research Laboratory at Los Alamos National Laboratory, according to University of Wisconsin Soil Science Laboratory protocol^21^. Briefly, the homogenized samples were dried at 105°C to remove moisture until weights were stable. Then samples were placed in a 360°C furnace and baked for 2 hours. Weights were taken before and after all steps to quantify moisture content and determine the loss-on-ignition (LOI). Testing of pH was performed by suspending soil in deionized (DI) water at a 1:2 (w:v) ratio. Measurements were taken with a SevenMulti pH Meter (Mettler Toledo, Columbus, OH).

For dissolved organic carbon (DOC) and dissolved total nitrogen (DTN) measurements, 1g of soil was added to 9 mL of sterilized DI water and vortexed. The solution was allowed to settle for 5 minutes, then the supernatant was filtered through a 0.22 µm Millex-GP PSE filter (Millipore Sigma) via syringe (Fisher Scientific, Hampton, NH, USA). Filtered extracts were analysed for non-purgeable organic carbon and total nitrogen using a Shimadzu TOC-LCSH/TNM-L Carbon/Nitrogen Analyzer (Shimadzu Scientific Instruments, Inc. Columbia, MD) employing a catalytic combustion oxidation method and non-dispersive infrared detector (NDIR) for carbon and chemiluminescent detector for nitrogen. Samples were acidified with 5N hydrochloric acid (HCl) and purged with air to remove inorganic carbon. A sample aliquot was then injected onto a palladium catalytic bed held at 720 °C. The CO2 and NO combustion products were then measured serially with the two detectors. The instrument was calibrated daily using a NIST-traceable standard of potassium hydrogen phthalate and potassium nitrate. Reagent water blanks were run periodically along with the check standards to monitor carry-over and instrument drift. During methods development a representative set of samples was analysed by the method of standard additions to verify complete recovery of organic carbon and total nitrogen under the analysis conditions.

### Genomics Preparation

DNA was extracted using the DNeasy PowerSoil Pro kit (Qiagen, Hilden, Germany). The concentration of DNA was obtained using the Qubit DNA Assay Kit (Thermo Fisher Scientific, San Jose, CA, USA). The quality of the DNA was determined by running the sample on an E-Gel 1% agarose gel (Thermo Fisher Scientific, San Jose, CA, USA) with Lambda DNA/HindIII Marker (Thermo Fisher Scientific, San Jose, CA, USA).

Illumina libraries were prepared using NEBNext Ultra DNA II Library Preparation Kit (New England Biolabs, Ipswich, MA, USA). 500 ng of DNA was fragmented using a Covaris E220, the ends made blunt, and adapters and indexes was added onto the ends of the fragments to generate Illumina libraries that can be sequenced on an Illumina sequencer. Illumina libraries are eluted in DNA Elution Buffer (Zymo Research, Irvine, CA, USA). The concentration of the amplicons pool was obtained using the Qubit dsDNA HS Assay (Thermo Fisher Scientific, San Jose, CA, USA). The average size of the library was determined by the Agilent High Sensitivity DNA Kit (Agilent, Santa Clara, CA, USA). An accurate library quantification was determined using the Library Quantification Kit – Illumina/Universal Kit (KAPA Biosystems, Wilmington, MA, USA).

Libraries were normalized to the same concentration based on the qPCR results. Each of the libraries were sequenced on the Illumina NextSeq 500 generating paired-end 151 bp reads using approximately 31% of the NextSeq 500 High-Output v2.5 Reagent Kit (300-cycle) (Illumina, San Diego, CA, USA).

DNA sequences were quality filtered using the “rqcfilter2” program from BBTools^22^. Quality-filtered DNA sequences were taxonomically classified using GOTTCHA2^23^. Microbial diversity analyses used the ‘vegan’ package in R (version 4.1.1)^24^.

### Proteomics Preparation

All three protein extraction methods started with the same mass of sieved soil. Three technical replicates of 250 mg each of arid soil were prepared for the initial extraction method comparison. Afterwards, protein abundance reproducibility was evaluated at three different soil masses (50 mg, 125 mg, and 250 mg), with five technical replicates for each mass group. The mass test samples were brought up to the same relative volume with buffer to ensure there was sufficient volume for the extraction methods. In both the protein extraction method comparison and mass comparison experiments, buffer was added in the final step of each extraction in a 2:1 (v/v) ratio to the soil to solubilize and lyse the extracted proteins. These samples then proceeded through the normal S-Trap™ protocol.

### Protein Extraction: Bead Beating with SDS Buffer

Sieved soil and the extraction buffer consisting of 5% SDS in 25 mM Tris-HCL (pH 8.0) were combined in a 2 mL bead beating tube containing 0.5 mm glass beads (Fisher Scientific). The tube contents were homogenized via vortexing for 10 minutes. Samples were filtered through a 0.22-µm Millex-GP PSE filter (Millipore Sigma) via syringe prior to digestion.

### Protein Extraction: Boiling with an SDS Buffer

Sieved soil and the extraction buffer consisting of 5% SDS in 25 mM Tris-HCL (pH 8.0) were combined in a 2 mL Eppendorf Lo-Bind tube (Eppendorf, Hamburg, Germany) and vortexed for 10 seconds to mix. The tubes were placed in a heating block (VWR, Radnor, PA, USA) preheated to 100 °C for 6 minutes. The tubes’ contents were allowed to cool to room temperature. Samples were filtered through a 0.22-µm Millex-GP PSE filter (Millipore Sigma) via syringe prior to digestion.

### Protein Extraction: Sodium Hydroxide Two-Phase Extraction

Sieved soil was placed into cryogenic tubes (Thermo Fisher Scientific, San Jose, CA, USA) with six 2.5 mm magnetized stainless-steel balls (McMaster-Carr, Elmhurst, IL, USA). Tubes were placed individually into a stainless-steel grinding tube without the center cylinder (Spex Sample Prep). Samples were milled under liquid nitrogen with a Spex Sample Prep 6875 high-capacity cryogenic grinder (Spex Sample Prep) for a 4-minute pre-cool followed by two 4-minute cycles at 25 Hz with a 4-minute cooling period in between cycles. After being allowed to return to room temperature, 10% polyvinylpolypyrrolidone (PVPP) (v:v) was added to cover the milled soil. Samples were mixed via vortexing for 30 seconds, allowed to sit for 10 minutes and then vortexed again for 30 seconds. Following this, 0.1 M sodium hydroxide (NaOH) was added at 1:1 (v/v) ratio of the soil, the samples were shaken at 150 rpm at 20 °C for 30 minutes with a thermo-mixer and centrifuged at 3,220 x g for 20 minutes before the supernatant was moved to a new tube and extracted 1:1 (v:v) with a biphasic 8:5 (v:v) phenol:water mix. The phenol phase was washed with equivalent amounts of fresh water, removing the water after each wash. 0.1 M ammonium acetate in methanol was added, creating a 5-fold dilution. The proteins were allowed to precipitate overnight at 4°C. After centrifuging at 10,640 x g for 20 minutes at 4°C, the supernatant was discarded, and the resulting pellet was re-suspended in SDS-lysis buffer prior to digestion.

### Protein Digestion

The Pierce™ BCA Protein Assay Kit (Thermo Fisher Scientific, San Jose, CA, USA) was used according to manufacturer’s protocol to determine protein content post extraction. Samples were normalized to 30 µg protein in 0.1 mL with LC-MS grade water (Thermo Fisher Scientific, San Jose, CA, USA) prior to digestion. S-Trap™ micro columns (Protifi) were used for protein digestion.

The procedure was performed according to the manufacturer’s protocol, replacing the 120 mM TCEP and 500 mM MMTS with 20 mM DTT and 40 mM IAA, respectively. Mass spectrometry-grade Trypsin/Lys-C mix (Promega) was used for digestion with an overnight incubation at 37°C. After digestion and elution, peptides were dried down in a LABCONCO sample concentrator.

### Analysis *via* High Resolution Mass Spectrometry

Prior to analysis, the peptides were resuspended in 30 µL of 0.1% formic acid in water. Samples were then injected (4 µL) on a NEO Vanquish nanoLC system paired with an Eclipse Tribrid mass spectrometer equipped with a FAIMS Pro interface (Thermo Fisher Scientific, San Jose, CA, USA). Peptide separation was performed on a EASYspray C18 column (50 cm x 75 um, Thermo Fisher Scientific, San Jose, CA) over a 65-minute gradient at a flow rate of 250 nL/min. Mobile phase A consisted of water with 0.1% formic acid and mobile phase B of acetonitrile with 0.1% formic acid. The LC gradient was 0-3.32 min: 5-10% B, 3.32-49.8 min: 10-35% B, 49.8-52.4 min: 35-70% B, 52.4-53.6 min: 70-90% B, 53.6-56.8 min: 90% hold B, 56.8-57.3 min: 90-5% B, 57.3-65.0 min: 5% hold B. FAIMS Pro was used at a constant -50 CV.

The soil mass tests were performed on a U3000 nanoLC system (Thermo Fisher Scientific, San Jose, CA, USA) paired with an Eclipse Tribrid mass spectrometer with the same column and flow settings without the FAIMS Duo source. Mass spectrometric data was acquired using data-dependent acquisition with 3s time cycles. Full MS scans were collected at 240,000 resolution over a m/z range of 375 to 1500. The top 20 monoisotopic peaks of most abundant peptides detected in each full scan with charge states 2–6 being isolated for MS2 analysis at normalized collision energy of 30. Fragmented ions were analyzed in the ion trap.

### Data Processing

De novo sequencing of raw data was performed with PEAKS Studio 11 (Bioinformatics Solutions, Waterloo, ON, CA). The output was filtered for a de novo score of ≥ 90. The data was filtered for duplicate peptides and a frequency of detection was determined for all peptides. The results were filtered to only include peptides that had a detection frequency of ≥5. These peptides were put into NIH BLAST with the Swiss-Prot database and filters Prokaryotae (taxid:2), Fungi (taxid: 4751), Viruses (taxid:10239), and Archaea (taxid: 2157) were applied. Percent identity was filtered for 90−100% and query coverage for 75−100%. All identifications were exported and filtered to find the organisms with the highest frequency, eliminating duplicates.

Libraries collected from the genomics preparation were sorted by relative abundance. The top ten species with the highest relative abundances across all replicates were identified. From these results, the top ten most abundant organisms were determined. FASTA files were found using UniProt Proteomes. Species without available proteomes were removed from the database list and the next most abundant species was promoted. Protein assignments were determined with Proteome Discoverer 3.0 (Thermo Fisher Scientific, San Jose, CA, USA) using Sequest HT and PSM Validator. Raw data from the mass test underwent protein assignment with Proteome Discoverer 3.0 using Sequest HT and PSM Validator. The genomically- and proteomically-informed databases determined in the protein extraction method comparison were used for the biased analysis.

## Results

### Soil Properties

Initial analyses performed on the soil included SOC, DOC, DTN, and pH. The results show that the soil is acidic with a pH of 5.28 and has low values of SOC, DOC and TN. The SOC was 1.8297 (g C/100 g soil) while the DOC and DTN were 0.0425 (mg C/g soil) and 0.0047 (mg N/g soil), respectively. The low carbon and nitrogen content in the soil is expected for surface soil from an arid shrub-grassland^25^.

### Reference Proteomic Database Building

Historically, microbiome proteomic studies relied solely on proteomic reference databases derived from metagenomics analyses (**Figure 1, red arrows**). We aimed to augment this approach with databases derived from soil protein extracts by using an approach known as de novo peptide sequencing (**Figure 1, blue arrows**). The complete workflow illustrated in Figure 1 visualizes all steps in the experimental design including sample preparation, analysis, and data processing. The metagenomic testing and database determination occurred in parallel with the proteomic sample preparation and determination of the proteomically-derived databases. The pipelines only overlap in the final database driven analysis. The top ten most abundant species for each pipeline were used in the final database driven analysis. This was done to allow for a faster protein assignment process and reduce the number of false identifications within the Proteome Discoverer software.

**Figure 1.**
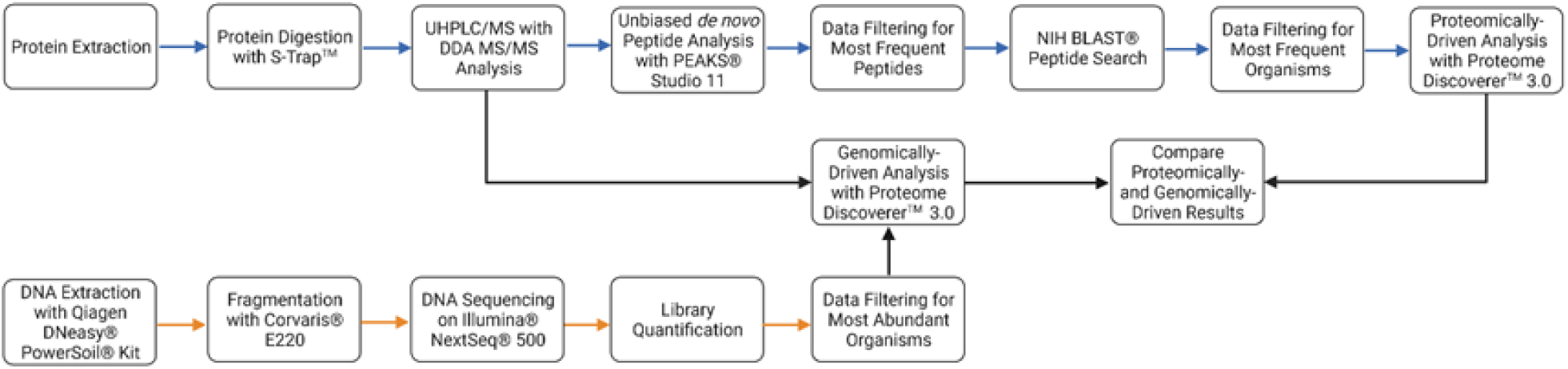
Workflow overview of the soil sample preparation and data analysis pipelines. The blue and orange arrows denote the proteomic and genomic pipelines, respectively. Black arrows represent where the pipelines overlap in the database driven analysis.

### Genomically-Derived Database Strategy

To establish the microbial diversity and composition, metagenomic analysis was performed for bacterial, archaeal, and viral species. In terms of relative abundance, Bacteria were dominant (87%) followed by Archaea (11%), and Viruses (1.5%). For bacteria, Actinobacteria (62%) was the most abundant followed by Chloroflexi (9.1%) and Bacteroidetes (8.9%). Euryarcheaota was the only phylum for Archaea (11%), which was composed of Methanomicrobia (7.8%) and Methanobacteria (3.2%) classes. For fungi, Ascomycota represented 95% relative abundance and Basidiomycota was the remaining 5%. The three classes of fungi in the arid soil were Eurotiomycetes (51%), Sordariomycetes (44%), and Agaricomycetes (5%). The genomic diversity of the arid soil at the genus level was low with a species richness of 206 microbial genera.

### Proteomically-Derived Database Strategy

Central to the metaproteomics strategy is protein extraction and digestion into peptides amenable for mass spectrometry-based sequencing. In this study, three protein extraction techniques were examined: 1) SDS boiling method, 2) SDS bead beating method, and 3) NaOH-based method. All extracts were digested into peptides using a modified S-Trap™ protocol and then de novo sequenced. The most frequently sequenced peptides, irrespective of extraction technique, were then associated with an organism using the NIH BLAST tool. The analysis resulted in 480,301 total identified peptides. There were 3,421 and 67 peptides remaining after filtering for a de novo score ≥90 and a frequency of ≥5, respectively. The most common peptide was sequenced 35 times. NIH BLAST searching yielded 597 organisms present. The proteomes with the highest and lowest number of identifications were 16 and 4. Organisms which had the most frequently sequenced peptides were rank ordered into a top ten list (**Table 1**). The ranking of these species is based on the frequency of occurrence across all the peptide identifications for arid soil.

**Table 1.**
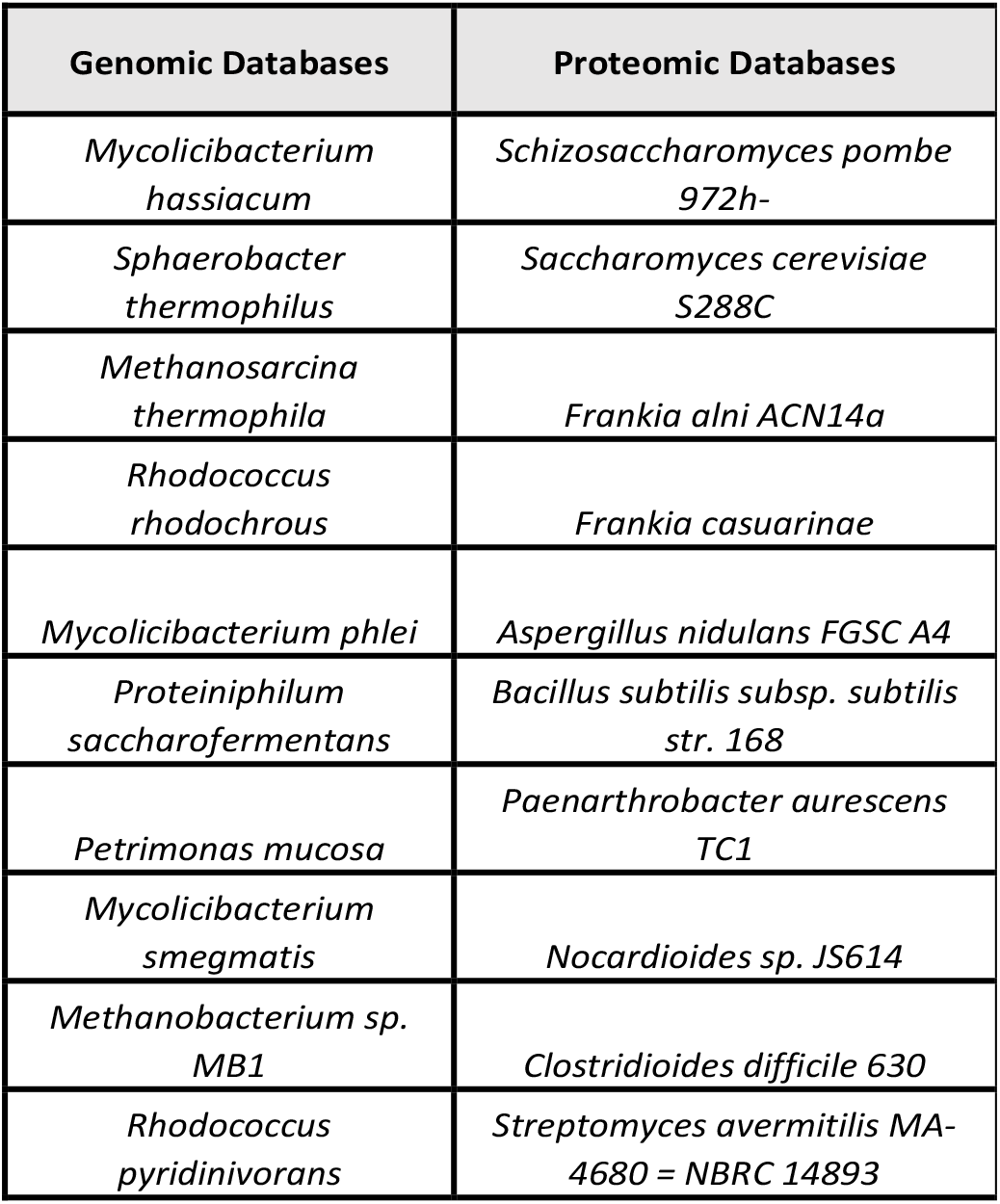
Genomically- and Proteomically-derived databases used for metaproteomics.

### Proteome Coverage Based upon Database Origin (DNA *vs*. Protein)

Using the top ten most abundant species from the metagenomic analysis, irrespective of extraction techniques, showed the arid soil to have 1,495 protein identifications. The species with the highest proportion of proteins identified was *Mycolicibacterium smegmatis* (235) (**Figure 2**). This species was ranked as the 8th most abundant species according to the initial metagenomic analysis. This is followed by *Rhodococcus rhodochrous* (188) and *Rhodococcus pyridinivorans* (166), which were 4th and 10th most abundant according the metagenomic testing, respectively (**Table 1**).

**Figure 2.**
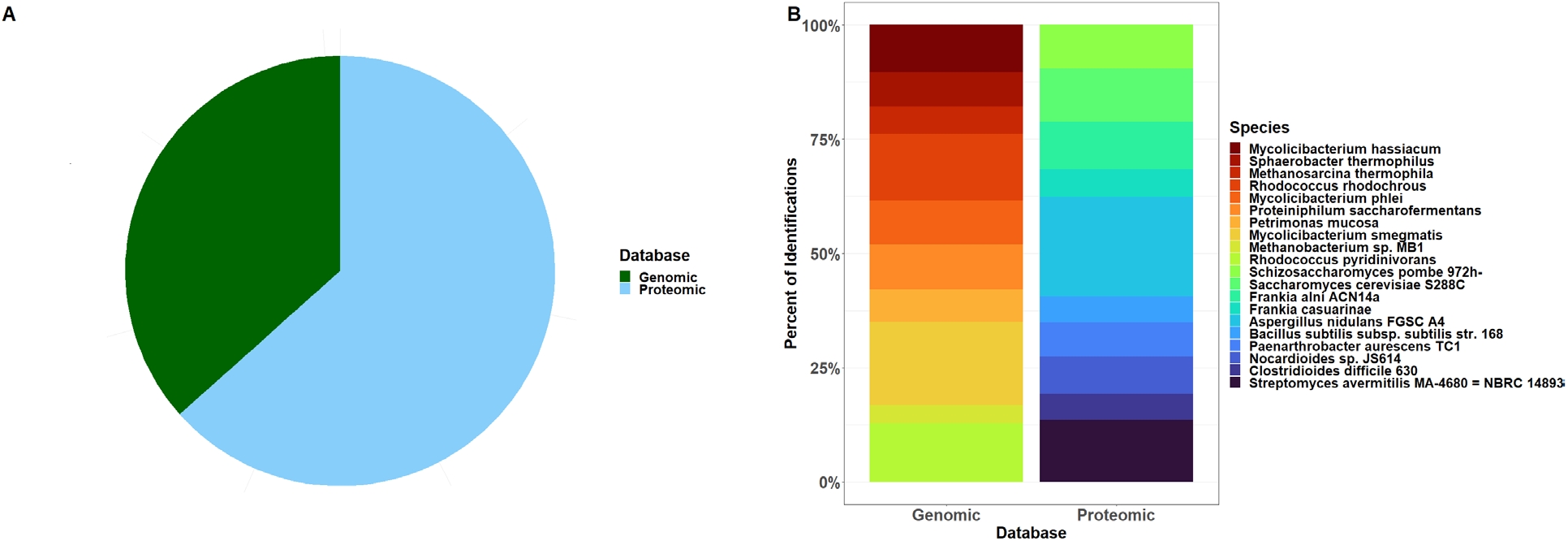
Contributions to the total proteome based upon database strategy for each soil type. A) Proportion of the percentage of total protein identifications for genomically- and proteomically-informed databases. B) Contribution to total proteome percentages for each of the top ten organisms identified in the soil upon database origin.

The *de novo* peptide analysis yielded 2,415 protein identifications with *Aspergillus nidulans FGSC A4* (487) having the highest proportion of protein identified despite being 5th in the number of occurrences from the NIH BLAST search (**Figure 2**). The 2nd and 3rd species with the most identified proteins were *Streptomyces avermitilis MA-4680 = NBRC 14893* (305) and *Saccharomyces cerevisiae S288C* (260) despite being ranked 10th and 2nd by the NIH BLAST search, respectively (**Table 1**). Combining the two strategies nearly doubled the number of protein identifications to 3,540 with the proteomically-driven pipeline accounting for a majority of the protein identifications made (61.8%).

### Assessment of Protein Extraction Techniques

Although the S-Trap™ protocol is typically used for biofluids, utilizing soil did not require significant modification to the protocols. As shown previously, this protocol has been used for solid or semi-solid material such as stool in order to assess protein content^26^. Importantly, the three extraction methods compared were able to provide sufficient protein content to be digested on the S-Trap™. Biased data analysis revealed the SDS buffer boiling method contained ∼ 12% of all identified proteins, the SDS buffer bead beating method ∼77%, and the NaOH-phenol extraction ∼15% (**Figure 3**). The SDS buffer bead beating method was determined to be the best method based on the number of protein identifications, and therefore this extraction technique was used for the soil mass testing that followed.

**Figure 3.**
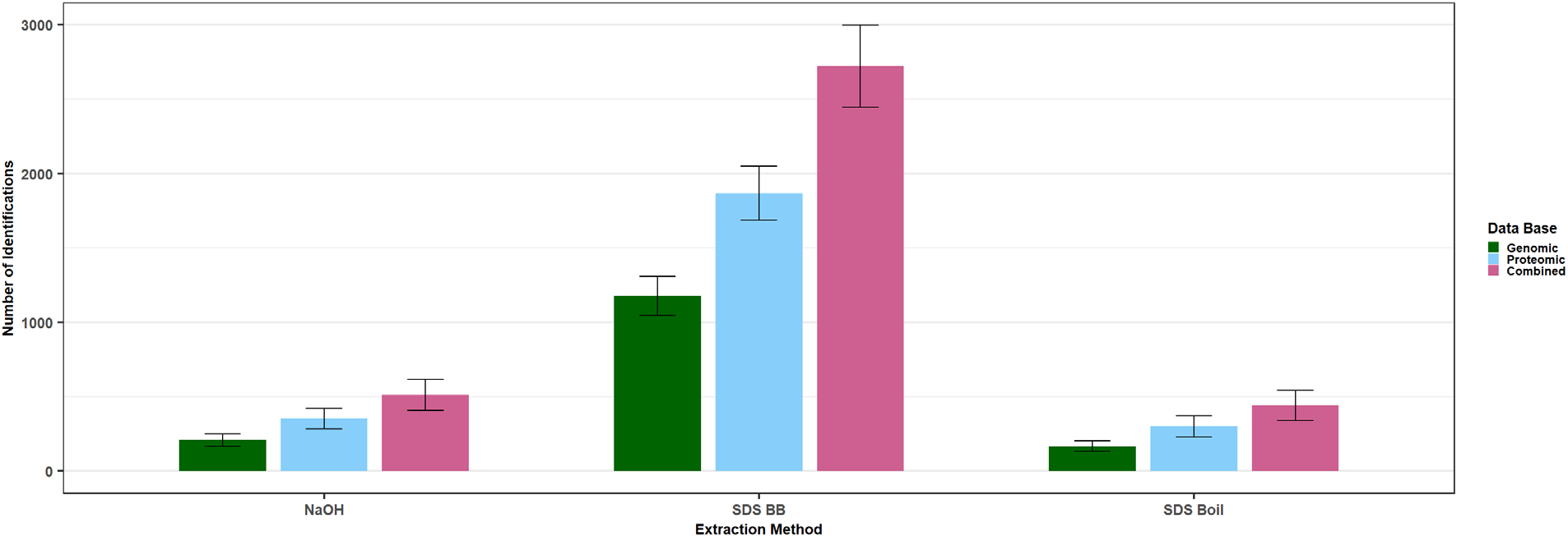
Assessment of proteome coverage based upon protein extraction method. Bar plot of the total number of protein identifications in soil for each extraction method separated by each database search set. Database set is denoted by color. The combined bar indicates the sum of both databases. The error bars represent the standard error of the mean (SEM) for each replicate.

### Assessing Minimal Amount of Soil Needed for Metaproteomics

Although soil analyses are often not limited by sample volume, the ability to increase analytical throughput is predicated upon being able to sample small soil volumes quickly and reproducibly. The S-Trap™ can be used in a 96-well format, which drastically increases throughput and allows for robotic sample manipulation. As shown in the protein extraction method comparison, the extraction method using the 5% SDS with Tris-HCl (pH 8.0) with bead beating yielded the highest number of protein identifications. This method was then applied with minute soil masses (≤250 mg) to ensure reproducible protein extraction is achievable from small sample sizes. Principal component analysis (PCA) shows significant grouping of the different mass groups with the greatest variation occurring in the smallest mass group (**Figure 4a**). Comparing total protein identifications, the bead beating method had a total of 18,839 identified proteins across all the databases with 5,852 being common across the three mass groups (**Figure 4b**). The largest proportion of protein identifications are also shared by at least two of the mass groups. The 50 mg mass group had the lowest protein content (12,307 protein identifications) and the 125 mg mass group had the highest protein content (14,220 protein identifications).

**Figure 4.**
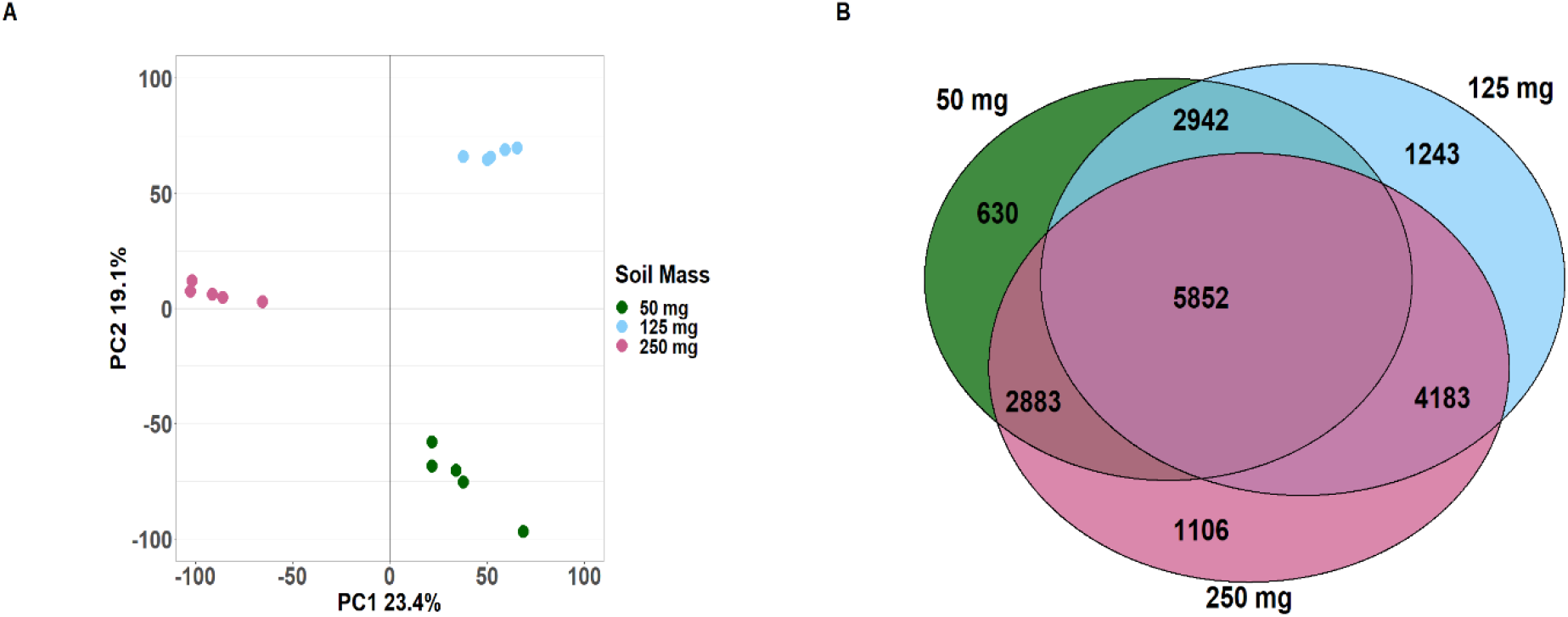
Proteome reproducibility and coverage based upon soil starting amount (grams). (A) PCA plot depicting the variability in protein assignments for the three mass groups [50, 125, 250 mg]. (B) Venn diagram shows the reproducibility in terms of total protein IDs based upon the amount of starting material.

**Figure 5.**
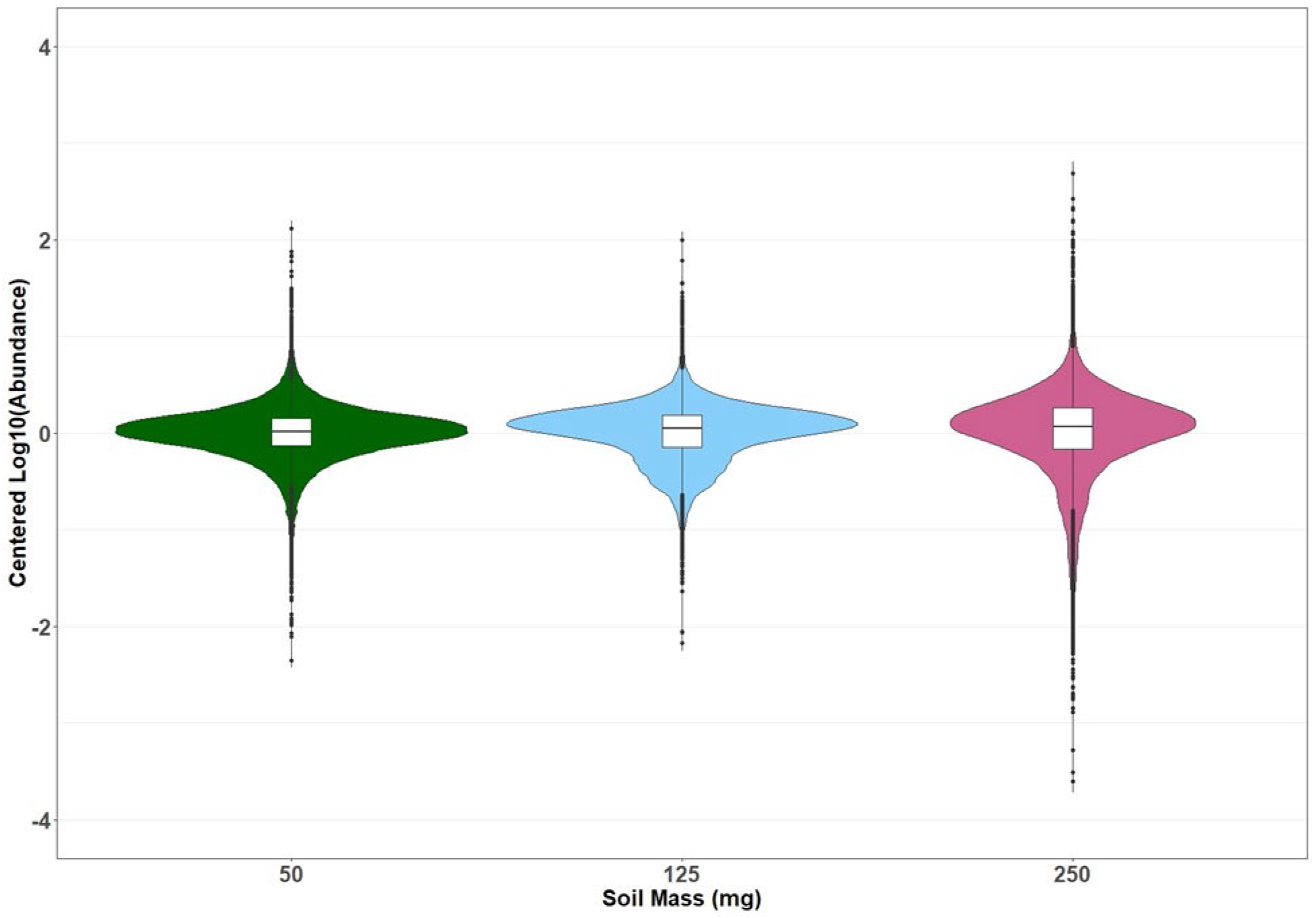
Replicate reproducibility of protein abundances within mass groups. Violin plot depicting the Centered Log10 of the abundance of the proteins common in all mass groups of all replicates of the three mass groups [50, 125, 250 mg]. Box plots were overlayed on the violin plots to better depict the median.

Assessing the precision in protein abundance between replicates was performed by comparing the abundances of the 5,852 proteins common across all mass groups. Replicates for each group were compared by centring the Log10 of the abundances against the grouped average abundance. The closer the centred abundance is to zero, the more consistent the replicates are with the group average. The Log10 of the abundances were taken prior to centering to negate the effect of outliers. The 50 mg and 125 mg mass groups show strong reproducibility between replicates with most of the points falling between -1 and 1 and the median centred close to zero. Like the other mass groups, a majority of the abundances from the 250 mg mass group fall within the same range with a median close to zero. However, the spread between this range is more pronounced indicating the difference between replicates is more noticeable.

## Discussion

### Complementarity of Genomic and Proteomic Reference Database Strategies

The proteomes used for the biased analyses are given in order of highest relative abundance across the two reference database strategies (Table 1). These were the top ten most abundant species according to the metagenomic and metaproteomic pipelines that also contained proteomes in the UniProt database. The databases present for the genomically informed workflow accounted for approximately 50% of the genomic makeup (**Figure 2**) of the soil based on the relative intensity. It is difficult to infer the coverage of the proteomically-informed databases; the proteomically-derived databases were picked based on the species that were most frequently observed in the NIH BLAST output.

Surprisingly, there is essentially no overlap between the genomically- and proteomically-informed database lists. Despite the proteomically- and genomically-derived databases producing nearly the same number of protein IDs there were nearly 40% more species (597 *vs*. 435) identified from the BLAST search. Conversely, there were nearly twice as many protein IDs (5,329 *vs*. 2,415) derived proteomically. However, the proteomic list does contain fungal species. While fungal species were included in the metagenomic testing, the results showed a low relative abundance across all replicates. Taken together, these data strongly suggest that proteomic and genomic databases taken alone are not sufficient to describe the full diversity of the soil microbiome, but instead offer a far more powerful tool when used in tandem.

### Protein Identifications and Replicate Reproducibility

It was determined that the bead beating method with the SDS-Tris HCl buffer was the most effective method based upon the number of protein identifications across all database-driven analyses (**Figure 3**). The soils’ properties (hydrophobicity, pH, humic substance content, etc.) could be a contributing factor to how well each extraction method performed. The use of surfactant-based extractions which do not remove humic acid substances has previously been shown to provide adequate protein extraction and can be used for a variety of soils, but arid soils with less protein content naturally contain less humic acid substances as supported by the experimental soil properties^27^. Phenol extraction has also been shown to produce a similar number of protein identifications as shown here, although within a related study a more laborious protein extraction (FASP) was used rather than the S-Trap™ protocol used here^28^.

### Novel Workflow Adaptable for High Throughput Applications

The methods using the SDS-Tris HCl buffer as the protein extraction buffer gave the best results. While neither of the other methods came close to the amount of protein extracted with the bead beating method, other methods of agitation such as boiling could prove to be effective on other soil types. The choice of protein extraction method should be carefully considered based on the soil type being studied. Furthermore, the two methods in which organisms are identified for database driven analysis produced two drastically different databases to search from. Despite this, the proportions of protein IDs remained consistent between the different extraction methods for the database set.

The increase in protein identifications between the method comparison and mass testing (∼3,000 to ∼18,000 proteins) could be attributed to method optimization and a larger amount of available data from the increased replicates (5 vs. 3). A lack of overlap in the replicates could also explain this increase, as each replicate would introduce a significant increase in unique protein identifications. The mass test data indicates that although there is increased variability when minute volumes of soil (50 mg) are extracted, these volumes are still well below the typical volumes utilized for DNA and RNA extraction. Within the proteins common to all mass groups, the variability between replicates for each group was similar. The variability within the 250 mg mass group was most pronounced which may have been caused by the protein load being too high during analysis. By incorporating the S-Trap workflow with soil analysis, it becomes possible to quickly multiplex protein extraction and analysis using 96-well plates and robotic sample handling. Because of the microbial diversity and stratification within soil, the ability to test smaller samples with higher throughput within the different layers of soil will undoubtedly increase our ability moving forward to catalogue these microbiotas and their changes.

## Conclusions

While a ‘universal’ soil extraction method is still to be developed, the speed, versatility, and simplicity of the S-Trap™ workflow used in this study have shown considerable improvement over the state of the art for soil metaproteome analysis. Future research could focus on identifying the factors that contribute to the differences in protein extraction efficiency between different soil types and finding optimal methods for database selection that include both genomically- and proteomically-informed methods to maximize protein identification^29^. Including the use of robotic sample preparation in addition to the S-Trap™ protocol could also be of interest in applications where smaller volumes of soil per sample could lead to higher throughput and even faster analysis. Given the extreme heterogeneity found within the soil microbiome, higher throughput could lead to less ‘pooling’ of results from soil samples and a better cross-section of the microbiome present within closely related samples.

## Author Contributions

ASM, Formal Analysis, Investigation, Methodology, Visualization, Writing-Original Draft, Writing-Review & Editing; PMM, Supervision, Methodology, Writing-Review & Editing; MEK, Conceptualization, Formal Analysis, Funding Acquisition, Investigation, Methodology, Project Administration, Resources, Writing-Original Draft, Writing-Review & Editing; EMM, Project Administration, Supervision, Visualization, Writing-Original Draft, Writing-Review & Editing; TGG, Conceptualization, Project Administration, Resources, Supervision, Visualization, Writing-Original Draft, Writing-Review & Editing.

## Acknowledgements

The U.S. Department of Energy supported this work through the Los Alamos National Laboratory. Los Alamos National Laboratory is operated by Triad National Security, LLC, for the National Nuclear Security Administration of the U.S. Department of Energy (Contract No. 89233218CNA000001). This work was also supported by the U.S. Department of Energy Biological System Science Division, through a Science Focus Area Grant (2019SFAF255).

